# Elucidating Chiral Myosin-Induced Actin Dynamics: From Single Filament Behavior to Collective Structures

**DOI:** 10.1101/2025.03.26.645581

**Authors:** Takeshi Haraguchi, Kohei Yoshimura, Yasuhiro Inoue, Takuma Imi, Koyo Hasegawa, Taisei Nagai, Toshifumi Mori, Kenji Matsuno, Kohji Ito

**Affiliations:** Department of Biology, Graduate School of Science, Chiba University, Chiba 263-8522, Japan; Center of Quantum Life Science for Structural Therapeutics (cQUEST), Chiba University; Department of Micro Engineering, Kyoto University, Kyoto 615-8540, Japan; Department of Applied Molecular Chemistry, Institute for Materials Chemistry and Engineering, Kyushu University, Fukuoka 816-8580 Japan; Department of Interdisciplinary Engineering Sciences, Interdisciplinary Graduate School of Engineering Sciences, Kyushu University, Fukuoka 816-8580, Japan; Department of Biological Sciences, Graduate School of Science, Osaka University, Osaka, 560-0043, Japan; Molecular Chirality Research Center, Chiba University; Plant Molecular Science Center, Chiba University

**Keywords:** myosin, actin, motor protein, chirality

## Abstract

The myosin superfamily comprises over 70 classes, each with multiple subclasses, and shows substantial diversity in properties such as velocity, ATPase activity, duty ratio, and directionality. This functional diversity enables the specialized roles of each myosin in various organisms, organs, and cell types. Recent studies have revealed that certain myosins induce chiral curved motions of actin filaments. However, this newly identified property remains largely unexplored. Here, we investigated this chiral motion *in vitro* using *Chara corallina* myosin XI (*Cc*XI), which drives fast counterclockwise (CCW) movement of actin filaments. This chiral motion arises from asymmetric displacement at the filament’s leading tip, and its curvature depends on the surface density of myosin. Surprisingly, at near-physiological actin concentrations, actin filaments exhibiting chiral curved motion undergo collective motion, spontaneously forming a novel structure—termed the actin chiral ring (ACR)—that exhibits persistent CCW rotation. ACRs display remarkable stability, continuing to rotate at their formation site until ATP is depleted, while maintaining their structure even after rotation ceases. This stability is unprecedented among reported collective motions of cytoskeletal proteins driven by various motors. Our findings demonstrate that myosins with chiral activity can autonomously organize actin filaments into stable, chiral structures through collective motion, providing new insights into actin self-organization by unconventional myosins. This study advances our understanding of the diverse functional roles of unconventional myosins and introduces a new paradigm for cytoskeletal organization.

**Significance Statement:** Myosins are motor proteins that move along actin filaments and support various intracellular functions. Recently, some myosins have been found to drive actin filaments along chiral curved paths, but the mechanisms and significance of this behavior remain poorly understood. Here, we analyzed this activity and discovered that it not only drives chiral curved motion of single actin filaments, but also organizes them into stable, unidirectionally rotating ring structures through collective motion. These rings, termed actin chiral rings (ACRs), spontaneously emerge at near-physiological actin concentrations. Our findings uncover a previously unrecognized organizing principle of actin self-assembly driven by myosin with chiral activity and provide a new framework for understanding how cytoskeletal chirality is made.

## Introduction

Phylogenetic analyses of myosin motor domain (MD) sequences have revealed remarkable diversity within the myosin superfamily. To date, at least 79 distinct classes have been identified (1). These myosin classes and subclasses exhibit broad functional diversity in properties such as actin sliding velocity, actin-activated ATPase activity, duty ratio (the fraction of ATPase cycles during which the motor domain remains strongly bound to actin), and directionality of actin filament movement (2). Recently, a novel property has been reported: rat myosin IC and *Drosophila* myosin ID drive actin filaments in a counterclockwise (CCW) trajectory, resulting in chiral curved motions (3, 4). This chiral activity is particularly noteworthy because the motor domain of *Drosophila* myosin ID plays a critical role in establishing left–right (L–R) asymmetry at both the cellular and organ levels (4-7). However, the mechanistic link between the chiral activity of myosin ID and the emergence of cellular or organismal chirality remains poorly understood.

Motility characteristics of myosin isoforms—such as sliding velocity and directionality—have been extensively analyzed using *in vitro* motility assays. In this method, myosin molecules are randomly immobilized onto a coverslip, and fluorescently labeled actin filaments are introduced along with ATP to visualize their movements (8). Although the filaments move in various directions across the surface, their velocity and directionality are consistent with those observed for the corresponding myosin isoforms *in vivo*. Therefore, *in vitro* motility assays serve as a reliable method for quantifying the velocities and directionality of myosin-driven actin movements. These assays typically use fluorescent actin at a concentration of 5 × 10^−4^ mg/ml (9), which is approximately 1/1,000 of the intracellular actin concentration. Such low concentrations are employed to allow for the clear observation of individual actin filament trajectories.

Recently, a modified *in vitro* motility assay has been developed to enable the tracking of individual actin filaments under conditions that approximate intracellular actin concentrations. This improvement was achieved by supplementing the assay buffer with non-fluorescent actin at physiological levels, together with a low concentration of fluorescently labeled actin. Remarkably, under these dense actin conditions, actin filaments driven by skeletal muscle myosin II no longer exhibited random motions. Instead, they displayed collective behaviors characterized by coordinated movement. These include polar clusters, nematic streams, and swirling vortices, all of which feature parallel alignment of filaments along their longitudinal axes (10-18).

To date, most *in vitro* studies of collective filament motion have employed skeletal muscle myosin II (SkII), also known as conventional myosin (10-18). However, within muscle sarcomeres—where SkII is endogenously localized—both myosin and actin are fixed, and collective motion does not occur. In contrast, intracellular actin movements in plant cells and non-muscle animal cells are primarily driven by unconventional myosins. These movements often exhibit ordered filament orientation, suggesting that unconventional myosins may contribute to intracellular actin alignment through collective motion. Given that unconventional myosins differ in their motile properties from SkII, their collective dynamics are also likely to be distinct.

Collective motion of the active matter is strongly governed by the geometric shape and motility characteristics of its components. For rod-shaped active matter, such as those composed of elongated filaments, collective motion is predominantly driven by lateral interactions, resulting in alignment along the longitudinal axes. This phenomenon has been observed in the collective motion of actin filaments driven by skeletal muscle myosin II, where filaments behave as linear rods and exhibit motion aligned with their long axes, as described previously (10-18). Recently, increasing attention has been directed toward the collective motion of chiral active matter, in which the components either possess intrinsic chirality or exhibit chiral motion patterns(19, 20). FtsZ, a bacterial homolog of the eukaryotic protein tubulin, polymerizes into filaments. When these filaments are anchored to a lipid surface via FtsA, they exhibit a treadmill-like motion and adopt a chiral shape. At high concentration, these filaments self-organize into rotating chiral rings, a process proposed to arise from lateral interactions between filaments with chiral geometry (21, 22).

In this study, we demonstrate that the motor domain of *Chara corallina* myosin XI (*Cc*XI MD) induces counterclockwise (CCW) actin movements. This chiral activity resembles that reported for rat myosin IC (3)and *Drosophila* myosin ID (4). However, unlike these class I myosins, *Cc*XI MD exhibits fast motility (23-25), enabling detailed analysis of chiral motion within a shorter time window. We show that *Cc*XI MD, a myosin with chiral activity, possess a unique capacity to autonomously drive the formation of highly stable, actin chiral structures—structures we term actin chiral rings (ACRs)—through collective motion of actin filaments. This finding provides new insights into the mechanisms of actin filament self-organization and broadens our understanding of the functional diversity of unconventional myosins in cytoskeletal dynamics.

## Results

### Chiral Motion of Actin Filaments by *Cc*XI

In the standard *in vitro* motility assay (Fig. 1A), we found that *Cc*XI MD, a plant myosin XI previously highlighted as a fast myosin (23-25), drove actin filaments in a counterclockwise (CCW) circular path when viewed from the actin side (clockwise (CW) when viewed from the objective lens side) (Fig. 1B: *Cc*XI MD and Movie S1). Similar CCW movements have been previously reported for actin filaments driven by mouse myosin IC (3) and *Drosophila* myosin ID (4). The curvature of actin movements driven by *Cc*XI MD was 17.0 ± 5.5 deg/µm (Fig. 1C: *Cc*XI MD), which exceeds the values reported for mouse myosin IC and *Drosophila* myosin ID using a similar assay (standard *in vitro* motility assay on glass surfaces) (3, 4). In contrast, actin filaments driven by rabbit skeletal muscle myosin II (SkII)—a widely used reference myosin—followed nearly straight trajectories, with an average curvature of 0.47 ± 8.8 deg/µm (Fig. 1B and C: Rab SkII).

**Figure 1.**
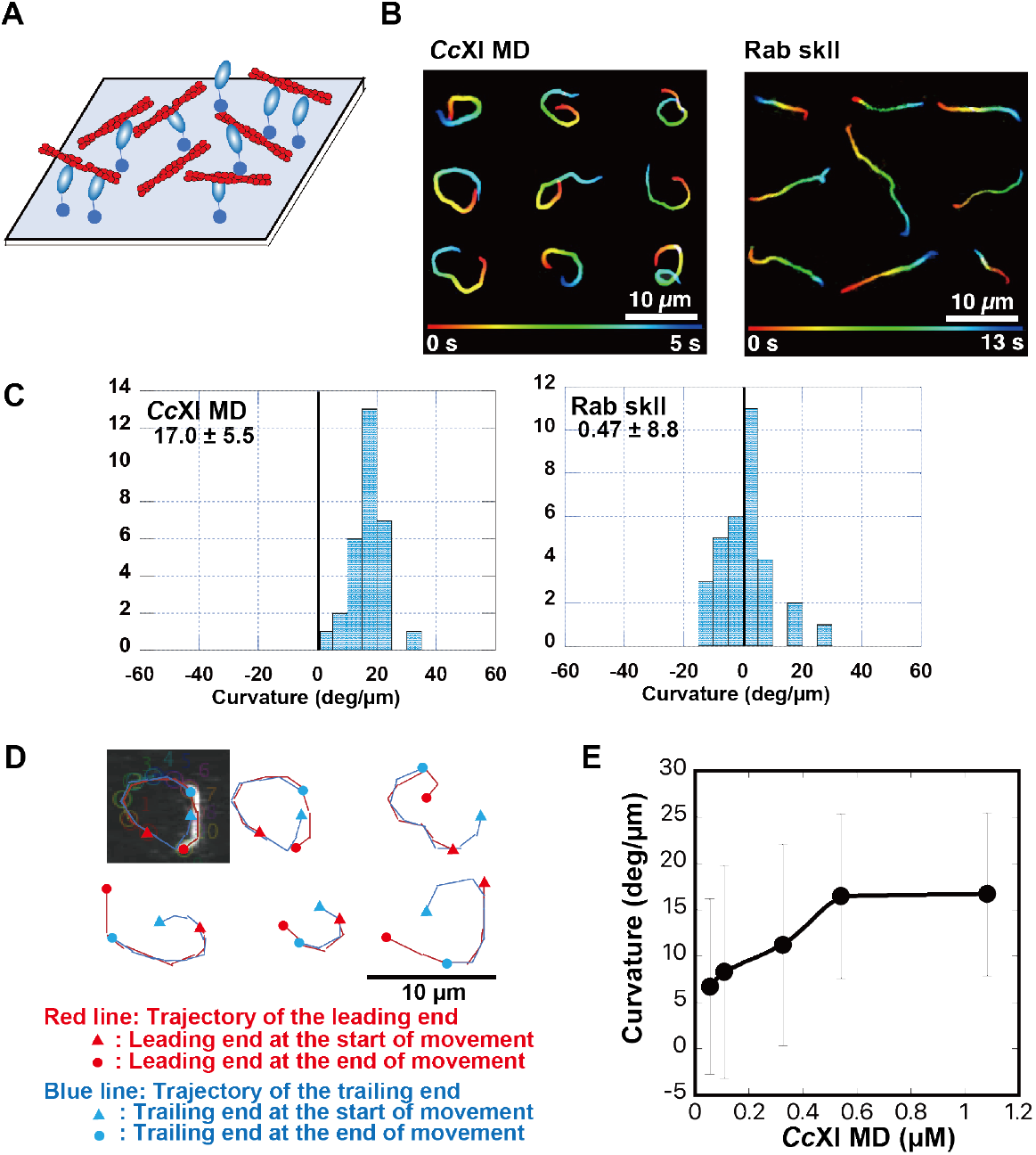
Chiral curved motion of actin filaments driven by *Cc*XI in the standard *in vitro* motility assay. (*A*) Schematic diagram of the standard *in vitro* motility assay. Myosin molecules were randomly immobilized on a coverslip, and fluorescently labeled actin filaments are added together with ATP to visualize filament motion. (*B*) Representative trajectories of actin filaments driven by *Chara corallina* myosin XI motor domain (*Cc*XI MD) and rabbit skeletal muscle myosin II (Rab SkII). For *Cc*XI MD, the trajectory represents filament motion over a 5-second interval, while for Rab SkII, the trajectory corresponds to a 13-second interval. In both cases, the color gradient from red to purple indicates the progression of time from the start to the end of the respective intervals. (*C*) Histogram showing the distribution of actin filament curvatures in the standard *in vitro* motility assay with *Cc*XI MD and Rab SkII. Curvature (degrees per micrometer) was calculated from orientation changes measured every 1 µm across 21 points along each filament. Positive values indicate CCW movement when viewed from the actin side. Each distribution represents measurements from 30 actin filaments per myosin condition (see *SI Appendix*, Supplementary Materials and Methods). (*D*) Overlay of trajectories of the leading (red) and trailing (blue) ends of a single actin filament in the standard *in vitro* motility assay with *Cc*XI MD. The trailing end follows the same path as the leading end, indicating that curvature is generated primarily at the filament tip. (*E*) Dependence of filament curvature on surface myosin concentration in the standard *in vitro* motility assay with *Cc*XI MD. The x-axis represents the myosin concentration (μM) applied to a chamber with a spacer thickness of 0.24 mm. The y-axis shows the mean filament curvature ± standard deviation (n = 27–30 filaments per condition).

To determine whether this chiral activity is a common feature among class XI myosins, we measured the curvature of the actin trajectories using *Cb*XI-3 (*Chara braunii* myosin XI-3) MD, which belongs to the same class as *Cc*XI (26). Although *Cb*XI-3 is a paralog of *Cc*XI and shares high sequence similarity (98%) and identity (88%) with *Cc*XI, the curvature of the actin trajectory induced by *Cb*XI-3 MD (3.1 ± 8.7 deg/µm) was significantly smaller than that induced by *Cc*XI MD. These results suggest that the curvature of the trajectory is determined by subtle variations in the myosin amino acid sequence, even among class XI myosins. This finding is consistent with previous reports indicating that only a subset of class I myosins generate curved actin trajectories whereas most do not (3, 4).

### Mechanistic Features of Chiral Motion

To elucidate the mechanism underlying this chiral curved motion, we tracked both the leading and trailing ends of individual actin filaments. We found that both ends followed the same trajectory (Fig. 1D), indicating that the curved motion is primarily generated by asymmetric displacement at the filament’s leading tip, while the remainder of the filament passively follows the path without actively contributing to the curvature. This observation suggests that the leading tip of the filament is uniquely permissive to curvature, potentially due to localized flexibility, as further discussed in the Discussion and *SI Appendix*, Fig. S1.

To investigate the dependence of actin filament curvature on myosin density, we performed a series of *in vitro* motility assays. In these assays, *Cc*XI MD was immobilized onto the coverslip surface in chambers with a spacer thickness of 0.24 mm, and the concentration of myosin introduced into the chamber was systematically varied. As the myosin concentration increased, a progressive increase in actin filament curvature was observed, reaching a plateau at 0.5 μM (Fig. 1E). At this concentration, the surface myosin density was estimated to be approximately 3.6 molecules per 10 × 10 nm^2^ area. Given that the diameter of the myosin XI MD is approximately 4 nm (26), this density likely corresponds to near-saturation of the coverslip surface. Above this concentration, no further increase in curvature was detected, suggesting that the effect plateaued due to saturation of available myosin-binding sites. These results indicate that actin filament curvature increases in proportion to surface myosin density until saturation is reached The potential mechanism underlying this density-dependent increase in curvature is further discussed in the Discussion and *SI Appendix*, Fig. S2.

### Self-Organization of Actin Chiral Rings (ACRs)

Traditionally, standard *in vitro* motility assays have been conducted using actin concentrations far below physiological levels, typically around 0.0005 mg/ml (9). These low concentrations are preferred because higher concentrations of fluorescently labeled actin can obscure individual filaments, making their movement difficult to observe. However, recent advances have enabled assays to be performed at actin concentrations closer to physiological levels (∼1 mg/ml) by combining a low concentration of fluorescently labeled actin with non-fluorescently labeled actin at physiological levels. Using this modified high-concentration approach, studies with skeletal muscle myosin II have revealed that collective motion can emerge even when myosin and actin filaments are randomly distributed on the coverslip. Remarkably, under these conditions, actin filaments exhibited highly organized collective motion, characterized by parallel filament arrays. Moreover, most filaments within these aligned arrays moved in the same direction, suggesting that collective motion not only facilitates parallel alignment of actin filaments but also promotes directional uniformity (10-18).

Given that skeletal muscle myosin II drives actin filaments along straight trajectories parallel to their longitudinal axes, whereas *Cc*XI induces chiral curved motion, we hypothesized that applying the modified motility assay with *Cc*XI under actin concentrations approximating physiological conditions might result in a different type of collective motion. To test this hypothesis, we performed the modified motility assay using *Cc*XI MD. We found that collective motion emerged effectively at initial actin concentrations between 0.1 and 0.3 mg/ml, with the strongest effects observed at 0.1 mg/ml. Therefore, subsequent experiments were conducted under this condition to maximize the emergence of collective behavior.

To perform the modified *in vitro* motility assay, we first introduced a buffer containing 0.1 mg/ml actin (without ATP) into a chamber precoated with *Cc*XI MD. After a 10-min incubation, an ATP-containing assay buffer (without actin) was introduced to remove unbound filaments. Under these conditions, the actin filament density reached approximately 15 ± 4.1 filaments/µm^2^ (*SI Appendix*, Fig. S3). This value is remarkably close to the reported threshold density (∼20 filaments/µm^2^) required for the emergence of collective motion in assays using skeletal muscle myosin II (11). The detailed procedure for this modified assay, including filament length control and fluorescent labeling strategy, is described in the Materials and Methods section.

Immediately after the ATP-containing buffer was added to the chamber, the *Cc*XI-driven actin filaments moved randomly. However, after approximately 20 minutes, they self-organized into a ring-like structure, in which the entire ring exhibited collective motion, rotating predominantly in the CCW direction when viewed from the actin side. Notably, this rotational direction matched the trajectories observed for individual actin filaments at lower actin concentrations. We designated this structure the actin chiral ring (ACR) (Fig. 2A, 2B, and Movie S2). The formation of ACRs was observed when either rhodamine-phalloidin-labeled or Cy3-labeled actin was used, indicating that ACR formation does not depend on the specific fluorescent label.

**Figure 2.**
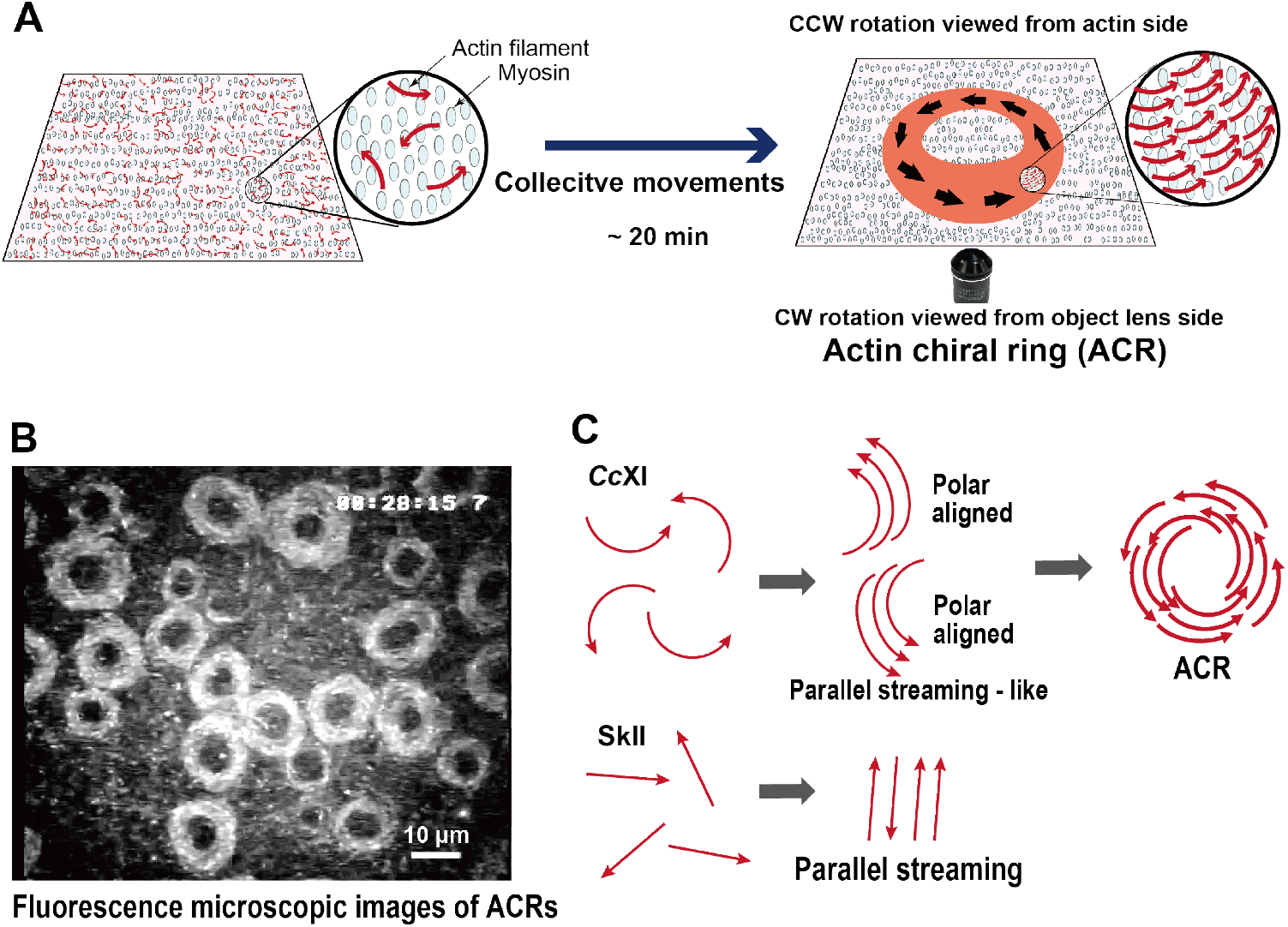
Autonomous formation of ACRs through the collective motion of chiral curved actin filaments driven by *Cc*XI in the modified *in vitro* motility assay. (*A*) Schematic illustration of ACR formation under elevated actin filament density. ACRs emerged within approximately 20 minutes after ATP addition, exhibiting CCW rotation when viewed from the actin side and CW rotation when viewed from the lens side. (*B*) Fluorescence microscopy images of fully formed ACRs. Once formed, the ACRs continued rotating in the CCW direction at the same location where they initially developed. The corresponding video is provided as Supplementary Movie 2. (*C*) Schematic representation of the formation process of collective actin motion under elevated actin filament density. ***Cc*XI**: Upon ATP addition, individual actin filaments driven by *Cc*XI initially exhibited CCW chiral curved motion and progressively aligned into a parallel, stream-like collective motion. These filaments then gradually assembled into ring-shaped structures, which subsequently rotated in the CCW direction. **SkII:** For comparison, a schematic of the collective motion of actin filaments driven by skeletal muscle myosin II (SkII), based on previous studies, is shown. In contrast to *Cc*XI, SkII drives straight actin filaments that form parallel streams without ring formation.

The dynamic process of ACR formation was continuously monitored over approximately 20 minutes, revealing distinct intermediate stages. To enable such continuous observation, prolonged exposure to excitation light was necessary, which partially compromised image clarity due to photobleaching. Nevertheless, key structural transitions were still clearly observable. Initially, actin filaments exhibited parallel, stream-like flow, which gradually coalesced into nascent ring-like structures. These nascent rings subsequently developed into fully formed ACRs, with their width increasing over time until reaching a stable state (Movie S3 and Fig. 2C: *Cc*XI).

### Structural Characterization of ACRs

We analyzed the structural characteristics of the ACRs. The ACRs exhibited limited size variation, indicating a high degree of structural uniformity. The outer and inner diameters were 21.8 ± 2.4 µm and 11.0 ± 2.3 µm, respectively (Fig. 3A). The inner curvature, calculated from the inner diameters, was 10.0 ± 2.3 deg/µm (Fig. 3A: IC). Although this value was lower than the curvature of individual actin filaments in the absence of collective motion (17.0 ± 5.5 deg/μm, Fig. 1C), the magnitude remained comparable, suggesting that a common underlying mechanism may contribute to ACR formation. This similarity, together with the consistent CCW direction of rotation, suggests that *Cc*XI-induced curved actin motion plays a critical role in ACR formation, although the curvature appears to be slightly attenuated during collective motion. In contrast, the outer curvature of the ACRs, calculated from the outer diameters, was 5.3 ± 0.6 deg/µm (Fig. 3A: OC), which was significantly lower than that of individual actin filaments in the absence of collective motion (17.0 ± 5.5 deg/μm, Fig. 1C). This reduced outer curvature likely imposes a physical constraint on the maximum size of the ACRs, contributing to their nearly uniform dimensions. Furthermore, the consistent size of ACRs below a certain threshold suggests that filament–filament affinity does not play a major role in their formation, as such a mechanism would be expected to result in continuous growth without a defined size limit.

**Figure 3.**
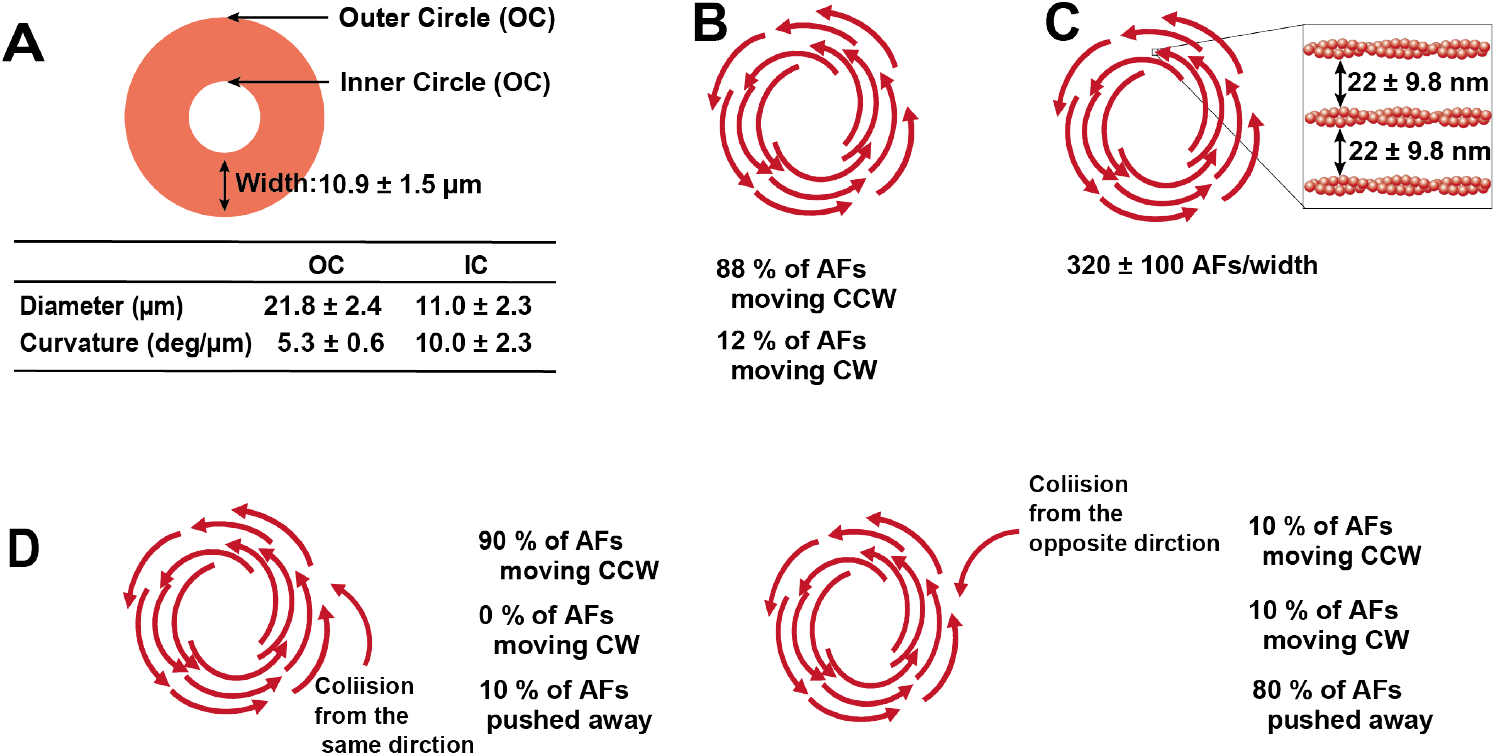
Analysis of actin filament motion within ACR structures. (*A*) Quantification of the out er and inner diameters of ACRs and the resulting ring width. The outer and inner diameters were determined by fitting circles in ImageJ. Data represent the mean ± standard deviation from 120 ACRs. Except for actin concentration, all assay conditions were identical to those used in the standard *in vitro* motility assay for measuring single-filament curvature, including a solution containing 25 mM Hepes-KOH (pH 7.4), 25 mM KCl, 4 mM MgCl_2_, 3 mM ATP, and 10 mM DTT. (*B*) Directional analysis of 90 actin filaments (AFs) from six ACR rings revealed that 79 AFs (88%) moved in the CCW direction, while 11 AFs (12%) moved in the CW direction. This analysis was performed by lowering the ratio of Cy3-labeled actin, as described in the SI *Appendix, Supplementary Materials and Methods*. (*C*) Estimation of filament density within ACRs. The labeling ratio was reduced to visualize individual filaments, as shown in *SI Appendix*, Fig. S5. Across 20 ACRs, the average number of filaments per ring width was 320 ± 20, with an interfilament spacing of 22 ± 9.8 nm, indicating relatively sparse filament packing within the ring. (*D*) Outcomes of filament-ACR collisions depending on the approach direction. (Left) When AFs approached in the same direction as ACR rotation, 90% were assimilated and adopted CCW rotation, while 10% were excluded. (Right) When AFs approached from the opposite direction, 80% were repelled, 10% were incorporated and rotated CCW, and the remaining 10% were incorporated but rotated CW. This latter population likely corresponds to the minority CW-rotating filaments observed in (B). Each category was quantified from observations of 30 AFs per approach direction.

The ACRs exhibited remarkable temporal and spatial stability. Once formed, they continued to rotate at the same location without positional displacement (Fig. 2B and Movie S2). When external fluid flow was intentionally applied after formation, the ACRs maintained their structure and continued to rotate at the same position, highlighting their structural integrity and robustness. Moreover, the ACRs preserved their ring-shaped architecture even after rotational motion ceased due to ATP depletion. Notably, ACRs were formed exclusively from nearly 100% pure actin and *Cc*XI MD (*SI Appendix*, Fig. S4), which were randomly distributed on the coverslip, indicating that neither actin-bundling proteins nor other contaminants were required for their formation. Taken together, these observations underscore the exceptional stability of the ACR as a structural unit composed solely of actin filaments.

To analyze the movement of individual actin filaments within the ACR, we adjusted the ratio of Cy3-labeled to non-fluorescent actin (*SI Appendix*, Supplementary Materials and Methods) to enable clear visualization of each filament. Our observations revealed that a substantial majority (88%) of actin filaments within the ACR rotated in the CCW direction, while a minority (12%) exhibited CW rotation (Fig. 3B). This result is somewhat surprising, given that the entire ACR appears to rotate in the CCW direction (Movie S2). This global rotation aligns with the directional bias of individual actin filaments driven by *CcX*I (Fig. 1B and C). The CW-rotating filaments likely represent those that entered the ACR from the opposite direction and continued to move without reorienting, as discussed below. Notably, the observation that 88% of filaments within the ACR exhibit CCW motion reflects a strong bias toward unidirectional alignment. This proportion is notably higher than the 60–80% directional alignment during the collective motion of parallel actin filaments induced by skeletal muscle myosin II (11). These results suggest that ACR formation facilitates more efficient unidirectional alignment of actin filaments compared to conventional collective motion systems.

To determine the number of actin filaments per ring width within the ACR, we employed the same approach of adjusting the ratio of Cy3-labeled to non-fluorescent actin (*SI Appendix*, Fig. S5). This method allowed us to estimate an average of 320 ± 10 filaments per ring width, corresponding to an interfilament spacing of approximately 22 ± 9.8 nm (Fig. 3C). This spacing—approximately three times the diameter of a single actin filament—suggests that filaments within the ACR are relatively spaced rather than densely packed.

To investigate the dynamics of chiral curved actin filaments interacting with the ACR, we again employed the method of adjusting the ratio of Cy3-labeled to non-fluorescent actin. This approach enabled detailed observation of collision outcomes. When actin filaments approached the ACR from the direction opposite to its rotation, 80% were repelled, 10% were incorporated into the ACR and began rotating in the CCW direction, and the remaining 10% were also incorporated but rotated in the CW direction (Fig. 3D, right). This latter group likely represents the minority CW-rotating population observed within the ACR (Fig. 3B, 12%). In contrast, when actin filaments approached from the same direction as the ACR’s rotation, 90% were incorporated and adopted CCW rotation, while the remaining 10% were excluded (Fig. 3D, left).

### Modulation of ACR Formation by Bundling Factors

In the cellular environment, actin-binding proteins play a crucial role in modulating actin networks by bundling actin filaments. To investigate the influence of actin-binding proteins on ACR formation, we focused on villin, a well-characterized actin-bundling protein. In modified *in vitro* motility assays using high actin concentrations, the presence of villin markedly accelerated ACR formation, reducing the required time from approximately 20 min (in the absence of villin) to just a few minutes (Fig. 4A, Movie S4). The ACRs formed in the presence of villin were significantly smaller, with an outer diameter of approximately 5 μm—about one quarter of the ∼20 μm diameter observed without villin. Notably, these smaller ACRs formed in the presence of villin maintained rotational behavior at their formation site (Fig. 4A, Movie S4), similar to those formed in the absence of villin. This observation suggests that villin-mediated size reduction does not impair either the rotational stability or the self-organization process of ACRs.

**Figure 4.**
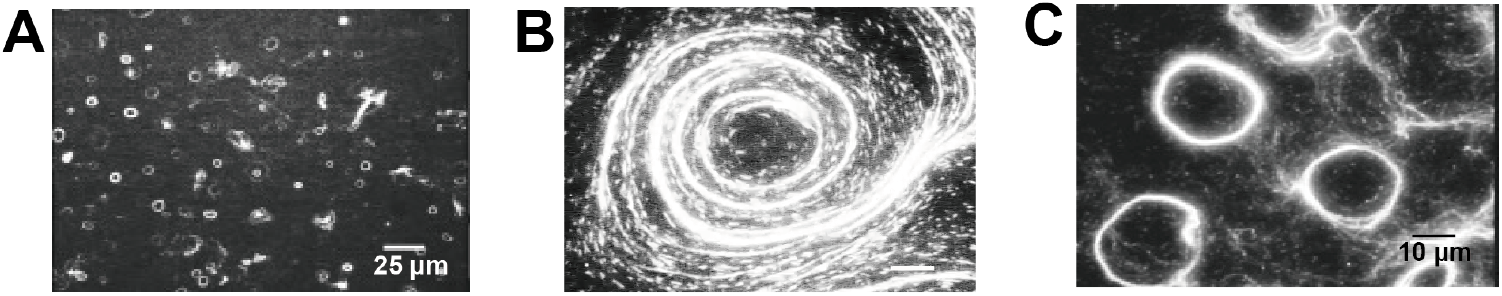
Effects of Actin-Bundling Protein and Methylcellulose on ACR Formation. (*A*) ACR formation in the presence of villin. To assess the impact of actin-bundling proteins on ACR formation, 1.5 µM villin was added to the modified *in vitro* motility assay. Within 1–2 minutes after the reaction began, numerous small ACRs (∼5 µm in outer diameter) appeared, rotating in the CCW direction. These smaller ACRs exhibited robust and continuous CCW rotation at their formation sites, closely resembling the behavior of ACRs formed in the absence of villin. The dynamic process is shown in Supplementary Movie 4. (*B*) ACR formation in the presence of methylcellulose. To mimic intracellular molecular crowding, 0.5% methylcellulose was included in the assay. Under this condition, large vortex-like ACRs (∼100 µm in diameter) emerged, rotating in the CCW direction. Unlike ACRs formed without methylcellulose, however, these structures were unstable and disintegrated shortly after formation. The corresponding dynamics are shown in Supplementary Movie 5. (*C*) ACR formation in the combined presence of villin and methylcellulose. When both 1.5 µM villin and 0.5% methylcellulose were added, ACRs with intermediate diameters (∼60 µm) formed. These ACRs also rotated in the CCW direction but, similar to those formed with methylcellulose alone, were unstable and disintegrated shortly after formation. The corresponding dynamics are shown in Supplementary Movie 6.

Methylcellulose, a polymer widely used in *in vitro* studies to mimic intracellular molecular crowding (27, 28), is known to promote actin bundling through the depletion effect (29, 30). Its presence markedly altered ACR formation, leading to the emergence of large, vortex-like ACRs with outer diameters reaching up to 100 μm—substantially larger than the ∼20 μm diameter observed in its absence. These vortex-like ACRs exhibited the same CCW rotation as those formed without methylcellulose (when viewed from the actin side). However, they were unstable and rapidly disintegrated after formation (Fig. 4B, Movie S5).

We next examined the combined effects of villin and methylcellulose on ACR formation. In the presence of both factors, ACRs exhibited slightly larger diameters—approximately 30 μm— compared to the typical ∼20 μm observed under standard conditions. These ACRs rotated in the CCW direction, consistent with the rotational direction observed in all other conditions. However, similar to the vortex-like ACRs formed in the presence of methylcellulose alone, they were less stable and more prone to disintegration (Fig. 4C, Movie S6).

These results suggest that while villin accelerates ACR formation and reduces its size, methylcellulose promotes the formation of larger and less stable ACR structures. Together, these two factors synergistically influence both the size and the stability of ACRs.

## Discussion

This study reveals a novel mode of actin self-organization driven by chiral myosin activity. We found that *Cc*XI, a fast plant myosin XI, drives actin filaments in the counterclockwise (CCW) direction. At actin concentrations comparable to those found in cells, this activity induces the autonomous formation of cell-sized actin chiral rings (ACRs) through collective motion. Remarkably, ACRs are exceptionally stable, maintaining continuous rotation until ATP is depleted, and preserving their structural integrity even thereafter.

### Mechanism of Myosin-Driven Chiral Actin Motion

While most myosins move actin filaments along the filament axis, certain class I myosins have been shown to induce chiral CCW motion (3, 4). However, detailed analyses of the motile behavior underlying this chiral motion have not yet been conducted, largely due to the extremely slow motility of class I myosins. In contrast, *Cc*XI induces rapid actin filament movement, allowing detailed analysis of chiral motion.

Our analysis reveals that the trailing end of the filament closely follows the curved trajectory of the leading end, indicating that chiral curvature arises primarily at the leading tip (Fig. 1D). We propose that this curvature results from an interaction between *Cc*XI and the intrinsic mechanical asymmetry of the actin filament. Specifically, the structurally flexible leading tip is susceptible to deflection by oblique forces generated by *Cc*XI power stroke, whereas the remainder of the filament is mechanically stabilized by multiple rigor-state attachments. This difference in structural flexibility— as illustrated in *SI Appendix*, Fig. S1—explains why curvature is localized at the leading tip. Moreover, we observed that curvature increases with surface myosin density (Fig. 1E). This finding contrasts with the well-established inverse relationship between actin velocity and myosin density, in which increased myosin density typically reduces actin filament velocity. This velocity reduction is thought to result from strongly bound myosin molecules hindering the progression of filaments driven by other myosins performing power strokes (31, 32). In contrast, the observed increase in actin filament curvature with higher myosin density likely originates at the filament’s leading tip, where elevated myosin density induces more frequent bending events—as illustrated in *SI Appendix*, Fig. S2—revealing a mechanistic distinction between curvature and velocity responses.

What drives the chiral curved motion of the filament’s leading tip? The widely accepted lever-arm model posits that myosin moves actin filaments via a swing of its lever arm during the power stroke (33). If the lever arm of *Cc*XI swings at an oblique angle relative to the filament axis, this could account for the observed curvature. To explore this possibility, we compared the rigor-state structures of acto-*Cc*XI MD, resolved by cryo-electron microscopy (34), with that of acto-SkII S1, also determined by cryo-electron microscopy (35). Structural analysis revealed that the lever arm positions of *Cc*XI and SkII in the rigor state were nearly identical, both aligning parallel to the actin filament (*SI Appendix*, Fig. S6). Thus, the post-power stroke conformation cannot account for the chiral curved motion. On the other hand, recent cryo-EM analysis of mouse myosin IC provides valuable insight into how chiral force generation might arise. That study demonstrated that the lever arm of myosin IC is tilted in the ADP-bound (pre-power stroke) state but becomes parallel in the rigor state, suggesting that an oblique power stroke originates from a pre-power stroke conformation (36). By analogy, *Cc*XI may also adopt an oblique lever arm orientation in the pre-power stroke state, generating the directional bending force responsible for chiral curved motion.

### Mechanism of ACR Formation

Under high actin concentrations, actin filaments undergoing chiral curved motion driven by *Cc*XI self-organized into ACRs through collective motion (Fig. 2A and B, Movies S2 and S3). To understand the mechanism behind ACR formation, it is informative to consider prior studies on actin self-organization driven by skeletal muscle myosin II (SkII). Under similar high actin concentrations, SkII promotes the emergence of ordered structures such as polar clusters, nematic streams, and vortices (10-18). In these systems, actin filaments align in parallel due to their liquid crystalline properties: at high densities, straight filaments behave like rod-shaped macromolecules that spontaneously align along their longitudinal axes to minimize free energy (Fig. 2C: SkII) (37, 38). In light of this framework and given the different interaction dynamics between straight and chiral curved filaments, it is unsurprising that *Cc*XI-driven collective motion results in a distinct structure. Based on the same liquid crystalline framework, we propose that chiral curved actin filaments initially align with the same polarity and subsequently accumulate to form ACRs (Fig. 2C: *Cc*XI). This two-step model is supported by time-lapse imaging (Movie S3), in which filament behavior closely mirrors the schematic sequence shown in Fig. 2C.

Previous studies suggested that parallel alignment of actin filaments under SkII arises from changes in filament orientation upon collision. However, two-filament collisions alone are insufficient to explain such alignment, implicating the necessity of multifilament interactions (12, 13). Consistent with this, we also observed that collisions between two chiral curved actin filaments rarely resulted in substantial angle changes, suggesting that multifilament interactions are likewise essential for effective alignment in chiral systems. To validate this idea, we simulated multifilament collisions using a simplified model in which each filament’s leading edge adjusts its orientation based on nearby filaments. For straight filaments, the model yielded bidirectional nematic alignment (Movie S7). In contrast, chiral curved filaments spontaneously formed curved polar arrays that self-organized into ACR-like structures (Movie S8), closely resembling experimental observations (Movie S3). Notably, unlike the straight filament simulations, which failed to generate polar alignment and remained nematically ordered, the chiral system produced ring structures with strong polar alignment. These results are consistent with the *in vitro* behavior of ACRs, in which 88% of actin filaments rotated in the same direction (Fig. 3B), whereas SkII-driven systems showed lower polar alignment of 60–80% (11).

These simulations further reflect our *in vitro* findings that *Cc*XI-driven filaments form more structurally uniform and stable assemblies than straight filaments. While SkII-induced systems generate a variety of patterns including waves and clusters, *Cc*XI consistently produced ACRs with uniform size and unidirectional CCW rotation. This high structural and directional coherence likely underpins the long-term stability of ACRs.

Interestingly, similar chiral rotating ring structures have been observed in the bacterial cytoskeletal system. FtsZ, a tubulin homolog, forms curved filaments that undergo chiral treadmilling when attached to lipid membranes via FtsA. At high densities, these filaments self-organize into rotating chiral rings *in vitro* (21, 22), and simulations support the autonomous formation of these structures (19, 22). Strikingly, this system also involves intrinsically chiral curved filaments that, through filament–filament interactions, assemble into rotating ring structures—a process remarkably similar to the ACR formation revealed in our study. Moreover, it is notable that, as with ACRs, computational models of FtsZ dynamics reproduce the spontaneous emergence of rotating chiral rings via collective self-organization. Together with the similarities observed in bacterial systems such as FtsZ, these findings raise the intriguing possibility that chiral ring formation through collective motion may represent a fundamental, evolutionarily conserved mechanism of cytoskeletal organization.

### Potential Roles of ACRs in Cell Dynamics

ACR-like structures—referred to as acquosomes in some reports—have been identified in various plant cell types, including *Arabidopsis*, tobacco BY2 cells, zinnia, and *Characean* algae (39-47). When cytoplasmic extracts from *Characean* algae were observed under a dark-field microscope, acquosome-like structures exhibited rotational motion on the glass surface in the same direction as the ACRs characterized in this study. Electron microscopy further showed that these structures consist of uniformly polarized actin filament bundles (43). These findings suggest that *in vivo* acquosomes share structural and dynamic properties with *in vitro* ACRs. Furthermore, the formation of acquosomes occurs transiently during specific cellular events, such as the onset of growth or response to injury, and depends on myosin activity (39-42, 44-47). In pollen tube cells, for example, acquosomes appear briefly before elongation begins and are subsequently reorganized into polarized actin bundles that guide tip growth (40, 41, 45). Such polarized bundles are essential for unidirectional cytoplasmic streaming, a fundamental mechanism for intracellular transport in plant cells. Previous *in vivo* studies have suggested that such structures can form autonomously within cells through interactions between actin filaments and myosin XI (48, 49), and that actin-bundling proteins such as plant villin facilitate their formation by crosslinking filaments with uniform polarity (50). In line with these in vivo observations, our *in vitro* findings closely mirror the behavior of acquosomes. ACRs, which arise from *Cc*XI-driven chiral actin motion, assemble into similarly polarized bundles (Fig. 2B and Supplementary Movies 2 and 3). Furthermore, villin significantly accelerates ACR formation *in vitro* (Fig. 4A, C and Supplementary Movies 4 and 6). Together, these results suggest that ACRs may serve as experimentally tractable *in vitro* analogs of acquosomes, thereby providing a useful model for understanding their organization and dynamics *in vivo*.

Recent studies have shown that many animal cells exhibit intrinsic chirality, characterized by left– right (L–R) asymmetry at the single-cell level. This cell chirality, often driven by cytoskeletal asymmetry—particularly in actin filaments—can propagate to affect multicellular alignment and contribute to asymmetric tissue morphogenesis. Notable examples include heart looping in vertebrates (51), gut and genitalia rotation in *Drosophila melanogaster* (7, 52, 53), and shell coiling direction in *Lymnaea stagnalis* (54). Among these examples, *Drosophila* myosin ID is particularly noteworthy. Similar to *Cc*XI, it induces chiral curved motion of actin filaments (4) and plays a critical role in establishing cellular and organ-level asymmetry in *Drosophila* (4-7). However, how myosin ID links molecular-scale chirality to mesoscopic cytoskeletal architecture remains poorly understood. In this context, ACRs—chiral cytoskeletal assemblies that emerge under physiological actin concentrations—may provide a useful framework. Given that ACRs are chiral cytoskeletal assemblies that emerge under physiological actin concentrations, it would be compelling to investigate whether myosin ID can also drive ACR formation *in vitro*, as observed with *Cc*XI. Such findings could illuminate how this myosin contributes to cell chirality. Moreover, exploring the potential existence of ACR-like structures in *Drosophila* cells in a myosin ID–dependent manner could further link cytoskeletal self-organization to developmental asymmetry.

### Unconventional Myosins with Chiral Activity

Although myosin function has been extensively studied, their potential chiral activity has received little attention—possibly leading to the oversight of myosins exhibiting this property. One reason for this may be that chiral activity is highly dependent on surface myosin density and requires sufficiently high concentrations, as demonstrated in Fig. 1E. However, conventional *in vitro* motility assays are typically conducted at low myosin densities to ensure smooth actin filament movement, thereby hindering the detection of chiral activity. As a result, despite numerous studies on *Cc*XI since 2003 (23-25, 34, 55, 56), its chiral activity has remained unidentified until the present study. It is therefore plausible that other myosins previously examined under low-density conditions may also possess latent chiral activity.

Uncovering the chiral activity of myosins opens new avenues for understanding how these motors regulate intracellular actin dynamics. Our findings demonstrate that myosin-driven chirality can promote the autonomous formation of stable and highly ordered actin architectures, adding a new dimension to cytoskeletal regulation. This discovery expands the functional repertoire of myosins beyond conventional parameters such as velocity, ATPase activity, duty ratio, and directionality. Further investigation into myosin chiral activity in physiological contexts—including its roles in cytoskeletal self-organization and cellular chirality—may provide deeper insight into how myosins coordinate cytoskeletal architecture and cell function.

## Materials and Methods

**(see also *SI Appendix, Supplementary Materials and Methods*)**

### Protein expression, and purification

*Cc*XI MD and *Cb*XI-3 MD were expressed in insect cells (High Five^TM^, Life Technologies) and purified using nickel-affinity and FLAG-affinity resins as previously described (25, 26). Further details are described in the *SI Appendix, Materials and Methods*. Skeletal muscle myosin II was purified from rabbits as described previously (57, 58). Skeletal muscle actin was purified from Chicken scapes using the method of Spudich and Watt (59). Cy3-labeled skeletal muscle actin was prepared as described previously (58, 60).

### Standard *in vitro* motility assay

The standard *in vitro* motility assay was performed as previously described (25, 26). Briefly, 0.2 mg/ml of anti-c-Myc antibody was perfused into the motility chamber. After washing with assay buffer, 0.15 mg/ml of myosin molecules with a c-Myc epitope tag were perfused into the chamber to immobilize the myosin molecules on the coverslip surface using an anti-c-Myc antibody. Next, the assay chamber was infused with either assay buffer containing 0.0003 mg/ml rhodamine-phalloidin-labeled actin or the same concentration of phalloidin-labeled Cy3-actin. The composition of the assay buffer was as follows: 25 mM Hepes-KOH (pH 7.4), 25 mM KCl, 4 mM MgCl2, 3 mM ATP, 10 mM DTT, and an oxygen scavenging system (120 µg/ml glucose oxidase, 12.8 mM glucose, and 20 µg/ml catalase). Observations of motility were conducted at 25°C.

### Modified *in vitro* motility assay using high actin concentration

Myosin molecules were immobilized on the coverslip using an anti-c-Myc antibody as described in the standard *in vitro* motility assay (24-26, 55, 61). The assay chamber was subsequently infused with assay buffer without ATP, either containing 0.005–0.01 mg/ml rhodamine-phalloidin-labeled actin and 0.1 mg/ml actin or assay buffer containing 0.005 mg/ml phalloidin-labeled Cy3-labeled actin and 0.1 mg/ml actin. The actin used was pre-treated with gelsolin (Sigma-Aldrich, product number: G8032) at a molar ratio of 1000:1 to actin in the presence of 1 mM CaCl_2_, resulting in an average length of approximately 5 µm.

After 10 minutes of incubation, the assay buffer with ATP was infused to remove unbound filaments and observation began. Cy3-labeled actin was prepared by labeling actin with Cy3-succinimide. To avoid motility artifacts, the proportion of Cy3-labeled actin molecules within the filaments was kept below 10%, as previously reported (58). The composition of the assay buffer was the same as that used for the standard *in vitro* motility assay: 25 mM Hepes-KOH (pH 7.4), 25 mM KCl, 4 mM MgCl_2_, 3 mM ATP, 10 mM DTT, and an oxygen scavenging system (120 µg/ml glucose oxidase, 12.8 mM glucose, and 20 µg/ml catalase). In some experiments, the concentration of KCl in the assay buffer was 150 mM. Observations of motility were conducted at 25°C.

Notably, ACRs also formed in the absence of gelsolin treatment. In such cases, longer actin filaments were initially present; however, over time, these long filaments were gradually severed during the *in vitro* motility assay, eventually reaching shorter lengths. As a result, the time required for ACR formation was slightly prolonged compared to that observed with gelsolin-treated actin. These observations suggest that efficient collective motion and ACR formation require actin filament lengths to be less than approximately 5 µm on average.

### Measuring the curvature of the trajectory of a single actin filament in the standard *in vitro* motility assay

An orientation difference-based method was used to calculate curvature from a set of points. Specifically, the curvature was calculated by dividing the orientation angle differences by the distance between two points using the orientation information of the point sequence (62). The X and Y coordinates of the leading end of a single actin filament were measured at 1 µm intervals over 21 points using Bohboh Soft (63). We then determined the curvature of the trajectory of a single actin filament from a set of 21 points spaced 1 µm apart by dividing the total angular variation by the total distance of movement across these 21 points. For each myosin type studied, curvature measurements were taken from approximately 30 actin filaments and then averaged to obtain the final values.

## Abbreviations

ACR: actin chiral ring
*Cc*XI: *Chara corallina* myosin XI
MD: motor domain
SkII: skeletal muscle myosin II
CCW: counterclockwise
CW: clockwise
*Cb*XI-3: *Chara braunii* myosin XI-3
AFs: actin filaments

## Acknowledgments

We thank Dr. Junichiro Yajima and Dr. Takayuki Nishizaka for their helpful discussions. We thank Dr. Shoji Baba and Dr. Kogiku Shiba for their support in using Bohboh Soft. We thank Dr. Yuichi Hiratsuka for helpful advice on the design of the color gradient used in Fig. 1B.We thank Dr. Naruki Sato and Mr. Shinryu Wakatsuki for providing rabbit skeletal muscle myosin II. RIKEN Arabidopsis full-length cDNA clone (pda11214) was provided by RIKEN BRC which is participating in the National BioResource Project of the MEXT/AMED, Japan. This work was supported by a Grant-in-Aid for Scientific Research (JP 22K20623, JP 24K09482 to T.H., JP 23K23303, JP 23KK0254, JP 24K21756 to T.M., JP 15H05863, JP 24H01284 to K.M., JP 23K05710, JP 22H04833, JP 20K06583, JP 17K07436 to K.I., JP 1715H01309 to K.I.) from the Japan Society for the Promotion of Science (JSPS), by ALCA (KI) from the Japan Science and Technology Agency (JST). The calculations were partially carried out at the Research Center for Computational Sciences in Okazaki (24-IMS-C105) and using MCRP-M at the Center for Computational Sciences, University of Tsukuba.

