## Supplementary Information for "Elucidating Chiral Myosin-Induced Actin Dynamics: From Single Filament Behavior to Collective Structures"

.

Takeshi Haraguchi, Kohei Yoshimura, Yasuhiro Inoue, Takuma Imi, Koyo Hasegawa, Taisei Nagai, Toshifumi Mori, Kenji Matsuno and Kohji Ito

Corresponding author: Kohji Ito


**This PDF file includes:**

Supplementary Materials and Methods
Figures S1 to S6
Legends for Movies S1 to S8
SI References

**Other supporting materials for this manuscript include the following:**

Movies S1 to S8

***Supplementary Materials and Methods***

**Protein engineering**

*Cc*XI MD

A baculovirus transfer vector for *Cc*XI MD (pFastBac *Cc*XI MD) was generated as follows: The cDNAs of motor domains of *Cc*XI (amino acid residues 1–746 of *Cc*XI) with optimized insect *Trichoplusia ni* codons were artificially synthesized by Eurofins Genomic. This was cut with BamHI and AgeI and legated with a BamHI–AgeI cut fragment of pFastBac *Cb*XI-1 MD (1). The resulting constructs pFastBac *Cc*XI MD encodes an N-terminal tag (MDYKDDDDKRS) containing the FLAG tag (DYKDDDDK), amino acid residues 1–746 of *Chara corallina* myosin XI heavy chain, a C-terminal tag (GGGEQKLISEEDLHHHHHHHHSRMDEKTTGWRGGHVVEGLAGELEQLRARLEHHPQGQREPSR) containing a flexible linker (GGG), a Myc-epitope sequence (EQKLISEEDL), a (His)8 tag and SBP tag (MDEKTTGWRGGHVVEGLAGELEQLRARLEHHPQGQREP). The amino acid sequence of this *Cb*XI-1 MD was the same as that used previously (2-4), except for the sequences of the N- and C-terminal tag regions.

*Cb*XI-3 MD

A baculovirus transfer vector for *Cb*XI-3 MD (pFastBac *Cb*XI-3 MD) was generated as previously shown (1). The resulting constructs, pFastBac *Cb*XI-3 MD encodes an N-terminal amino acids (MDYKDDDDKRS) containing the FLAG tag (DYKDDDDK), amino acid residues 1–747 of *Chara braunii* myosin XI-3, and C-terminal amino acids (GGGEQKLISEEDLHHHHHHHHSRMDEKTTGWRGGHVVEGLAGELEQLRARLEHHPQGQREPSR) containing a flexible linker (GGG), a Myc-epitope sequence (EQKLISEEDL), His tag (HHHHHHHH) and SBP tag (MDEKTTGWRGGHVVEGLAGELEQLRARLEHHPQGQREP).

Baculovirus transfer vectors of pFastBac *Cc*XI MD and *Cb*XI-3 MD were expressed in insect cells (High Five$^{TM}$, Life Technologies) and the expressed proteins were purified using nickle-affinity and FLAG-affinity resins as previously described (1, 4).

AtVillin 1

The full-length cDNA of *Arabidopsis thaliana* Villin 1 (*AtVillin 1*, AT2G29890.1) was provided by the RIKEN BRC through the National BioResource Project of MEXT/AMED, Japan (RIKEN Arabidopsis full-length cDNA clone, resource number: pda11214). The cDNA was subcloned into the pRSET expression vector (Invitrogen, Carlsbad, CA, USA), which contains an N-terminal His-tag. *Escherichia coli* strain BL21 (DE3) was transformed with the resulting expression plasmid. Transformed cells were cultured in LB broth supplemented with 50 μg/ml ampicillin at 20°C until the optical density at 600 nm (OD600) reached 0.6. Protein expression was induced by adding 0.4 mM IPTG, followed by incubation at 16°C for 16 hours. The *E. coli* cells expressing AtVillin 1 were harvested by centrifugation at 1,400 × *g* for 12 minutes at 4°C. The cell pellet was resuspended and lysed in lysis buffer containing 300 mM KCl, 0.1 mM EGTA, 10 mM β-mercaptoethanol (β-ME), 0.5% Triton X-100, 2 mg/ml lysozyme, 10 mM HEPES-KOH (pH 8.0), and a protease inhibitor cocktail (50 μg/ml leupeptin, 5 μg/ml pepstatin A, 0.25 mM PMSF, and 5 μg/ml chymostatin). The cells were flash-frozen in liquid nitrogen, thawed, and subjected to sonication to ensure complete lysis. The lysate was centrifuged at 38,000 × *g* for 30 minutes at 4°C. The resulting supernatant was incubated with nickel-nitrilotriacetic acid (Ni-NTA) agarose (Qiagen) on a rotating wheel for 1.5 hours at 4°C. The resin suspension was loaded onto a plastic column and washed with 200 ml of wash buffer containing 300 mM KCl, 10% glycerol, 15 mM imidazole, 10 mM β-ME, 10 mM HEPES-KOH (pH 8.0), and the same protease inhibitor cocktail. A second wash was performed using 15 ml of buffer containing 10% glycerol, 15 mM imidazole, 10 mM β-ME, and the protease inhibitor cocktail. AtVillin 1 was eluted with buffer containing 10% glycerol, 200 mM imidazole, 10 mM β-ME, and the same protease inhibitors.

**Analysis of movement direction of actin filaments within ACRs.**
To investigate the movement direction of actin filaments within the ACRs, we reduced the ratio of fluorescently labeled (Cy3) actin to non-fluorescent actin to 1:100. For comparison, in the modified *in vitro* motility assay using a high actin concentration for ACR observation, the typical labeling ratio was between 1:10 and 1:20 (see *Main Manuscript, Materials and Methods*).

**Simulation on motility assay of multiple actin filaments.**
To perform simulations on motility assay of multiple actin filaments, we construct a mathematical model to express the movement of the actin filament. In the model, each actin filament consists of $n_s$ segments of which length is $\sigma$. Assuming that the leading tip of the $i$-th actin filament could be followed by the rest of the segments, the movement of the filament can be described in terms of the leading tip motion using the following equation of motion (5):
$x_i(t+\Delta t) = x_i(t) + v_{\mathrm{a}}\Delta t\cos(\theta_i + \Delta\theta_i)$ (1)
$y_i(t+\Delta t) = y_i(t) + v_{\mathrm{a}}\Delta t\sin(\theta_i + \Delta\theta_i)$ (2)
where $x_i$ and $y_i$ are the $x$ and $y$ components of the position vector of the leading tip of the $i$-th actin filament, respectively. In the model, the sliding velocity of each actin filament is expressed by the average sliding velocity, $v_{\mathrm{a}}$, and the direction of the motion of the $i$-th actin filament is expressed by the angle $\theta_i$ measured counterclockwise from the $x$ axis on $x$ −y plane. The change of moving direction of the leading tip of the $i$-th actin filament, $\Delta\theta_i$, is calculated by the following equation:

$$\Delta\theta_i(t) = \frac{\delta^2 f_{\mathrm{myo}} v_{\mathrm{a}} \Delta t}{3k_{\mathrm{B}} T L_p} + \dot{\Theta}_{\mathrm{nem}}(\theta_i)\Delta t. \quad (3)$$

Here, $f_{\mathrm{myo}}$ is a myosin-induced active force normal to the filament axis at the tip of the $i$-th actin filament, δ is the mean interspace of myosin heads on the substrate, $k_{\mathrm{B}}$ is Boltzmann constant, $T$ is the temperature. $\dot{\Theta}_{\mathrm{nem}}(\theta_i)$ is a directional change rate of the tip per unit time through nematic interactions between actin filaments. To express $\dot{\Theta}_{\mathrm{nem}}(\theta_i)$ based on a filament alignment interaction, we refer to an energy function describing the intermolecular pairwise interaction of nematic liquid crystal (6). By differentiating this energy function with respect to the angle difference, $\dot{\Theta}_{\mathrm{nem}}(\theta_i)$ is derived as

$$\dot{\Theta}_{\mathrm{nem}}(\theta_i) = \frac{1}{\tau}\sum_j \cos(\theta_j - \theta_i)\sin(\theta_j - \theta_i) \quad (4)$$

where the relaxation time, $\tau$, is introduced to express kinetics of the filament alignment in our model. The summation in the right-hand side of Eq. (4) is applied over the $j$-th segment of a filament if the $j$-th segment of the filament is located within a cut-off distance of $\sigma/2$ from the the tip of the $i$-th filament, and the angle difference $|\theta_j - \theta_i|$ exceeds the nematic threshold angle $\theta_c$. Because we allow filaments to be overlapped in the simulations, we ignored the explicit contribution of the inverse of the distance between filaments such as a weight of interaction. The model parameters are listed in as mentioned below.

- Filament density: 12.5 filaments/μm²
- Filament length: 1 μm
- Filament velocity: 2 μm/s
- Myosin spacing: 0.005 μm/molecule
- Mean velocity of sliding motion of the filament: 2 μm/s
- Persistence length of the filament: 10 μm
- Length of the coarse-grained filament segment: 5 x $10^{-2}$ μm
- The number of segments of the filament: 19
- Temperature: 310 K
- Normal force density acting on the filament tip: -2 x $10^{-2}$ μN/μm
- Relaxation time for aligning filaments: 1 x $10^{-2}$ s
- Nematic threshold angle: 0.451 rad

Periodic boundaries are adopted for *x* and *y* directions, where the unit box size is $20 \times 20\ \mu\mathrm{m}^2$.

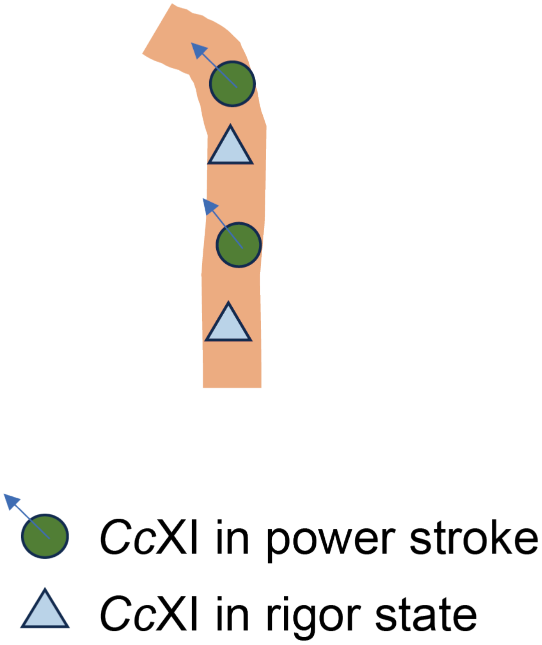


**Fig. S1. Hypothetical mechanism underlying curvature localized at the leading tip of the filament.** The leading tip of the filament, which exhibits relatively high structural flexibility, undergoes angular deflection in response to obliquely oriented forces relative to the filament axis, most likely generated by the power stroke of *Cc*XI. This localized bending arises because no rigor-state *Cc*XI molecules are bound to the anterior-facing side of the filament, allowing the leading tip to respond freely to directional forces. In contrast, the remainder of the filament is constrained by multiple rigor-state interactions with *Cc*XI at both anterior and posterior positions, effectively suppressing deformation in these regions. This spatial asymmetry in mechanical compliance accounts for the confinement of curvature to the leading tip, while the rest of the filament maintains structural rigidity.

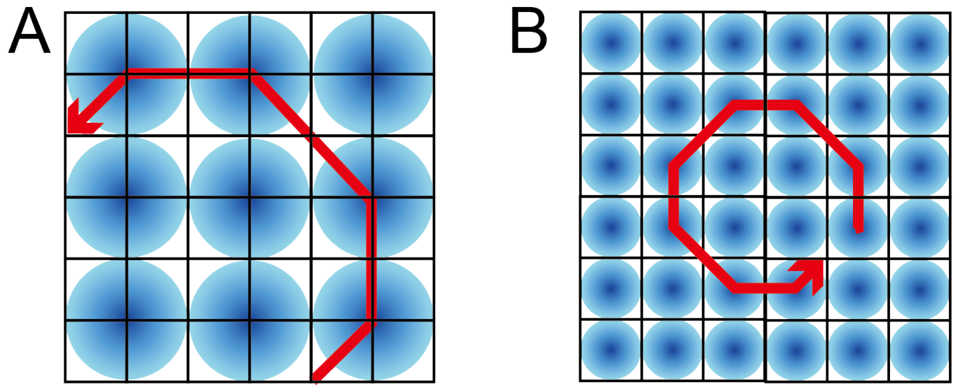


**Fig. S2. Model illustrating the dependence of actin filament curvature on myosin density.** In both panels, myosin motors are positioned at the center of circular areas. (A) Low-density condition: Nine myosin motors are distributed within the unit area, resulting in a relatively low myosin density. (B) High-density condition: Thirty-six myosin motors occupy the same unit area, increasing the myosin density by a factor of four compared to panel A. In both conditions, actin filaments interact with myosin at their leading tips, where oblique power strokes induce angular displacements of 45° in the counterclockwise (CCW) direction. Although the angular displacement per stroke is identical in both panels, the overall curvature of actin motion increases under high-density conditions, as evident from the comparison between panels A and B. This enhanced curvature arises from the increased frequency of tip deflections due to more frequent myosin encounters at higher surface density.

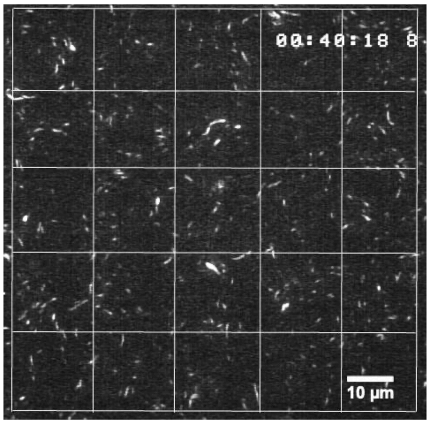


**Fig. S3. Estimation of actin filament density in the modified *in vitro* motility assay.**

To estimate the density of actin filaments during collective motion under high actin concentration, we conducted experiments using a reduced concentration of fluorescently labeled actin. Consistent with the conditions used for ACR observation, actin filaments were pre-treated with gelsolin to control their length. Specifically, 0.00033 mg/ml of Cy3-labeled actin and 0.1 mg/ml of non-fluorescent actin were used, yielding a labeling ratio of 1:300. This ratio is lower than the labeling ratio used in the ACR observation experiments (1:10–1:20; see Main Text, Materials and Methods). The number of Cy3-labeled actin filaments per unit area was counted immediately after the start of observation—prior to ACR formation—and this value was multiplied by 300 to estimate the total filament number. Observations were conducted across ten fields of view (90 μm × 90 μm), each divided into 25 squares (18 μm × 18 μm) for filament counting. In the representative field shown in this figure, 337 Cy3-labeled filaments were detected, corresponding to an estimated total of 337 × 300 = 101,100 filaments. This yields a filament density of 101,100 filaments / 8100 μm² = 12.5 filaments/μm². The average filament density across ten fields was 15.1 ± 4.1 filaments/μm². Scale bar = 10 μm

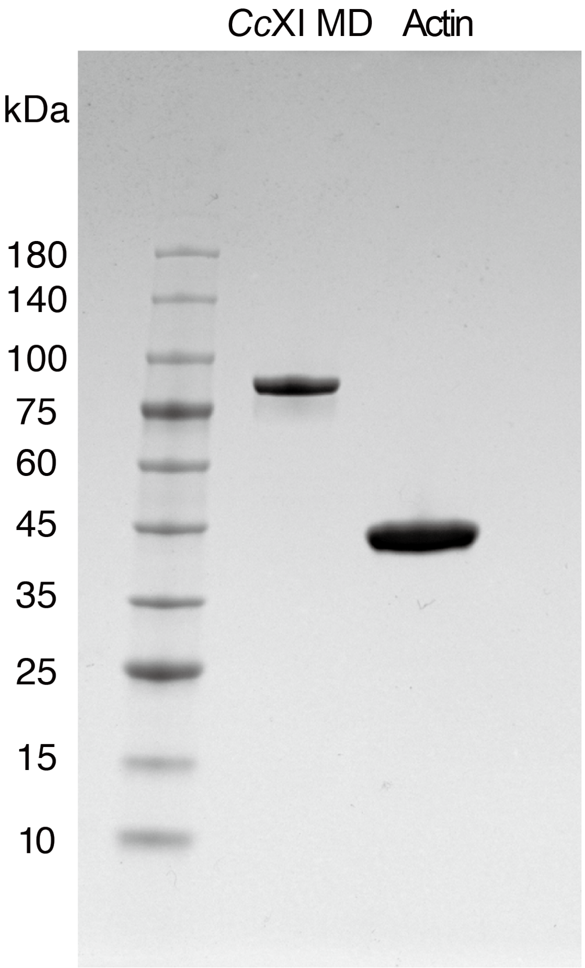


**Fig. S4. SDS-PAGE analysis of purified *Cc*XI MD and actin.**

Purified *Cc*XI MD and actin were analyzed by SDS-PAGE using a 4–20% polyacrylamide gradient gel and stained with Coomassie Brilliant Blue. Molecular mass markers (kDa) are indicated on the left. Both proteins displayed sharp and distinct bands, indicating near-homogeneous purity with no detectable contamination. These results confirm that the formation of actin chiral rings (ACRs) observed in this study was not attributable to the presence of other proteins, such as actin-bundling factors.

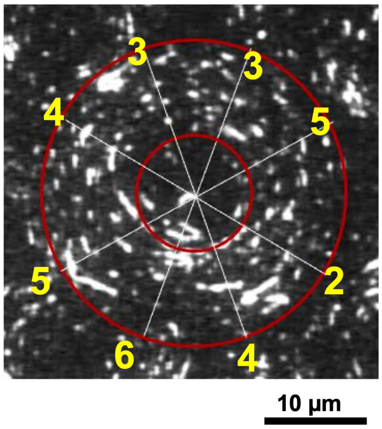


**Fig. S5. Estimation of the number of actin filaments per ring width within the ACR.**

To estimate the number of actin filaments spanning the ring width of an ACR, we used 0.001 mg/ml of Cy3-labeled actin together with 0.1 mg/ml of non-fluorescent actin, yielding a labeling ratio of 1:100. At a specific time point, four radial lines were drawn from the center of a representative ACR, generating eight radii. The number of Cy3-labeled actin filaments intersecting each radius within the defined ring width (i.e., the area between the inner and outer red circles) was counted. Each count was multiplied by 100 to estimate the total number of filaments, based on the labeling ratio. As shown in the figure, the yellow numbers represent the counts of labeled filaments per radius, with an average value of four. From this, the number of actin filaments within a ring width of 9 μm was estimated to be approximately 4 × 100 = 400. This corresponds to an estimated total of approximately 4 × 100 = 400 filaments within a ring width of 9 μm. Assuming an actin filament diameter of 8 nm, the inter-filament spacing was estimated to be approximately 15 nm. Fig. 3C summarizes data from 20 ACRs, showing an average of 320 ± 100 filaments per ring width, with an inter-filament spacing of 22 ± 9.8 nm (mean ± standard deviation).

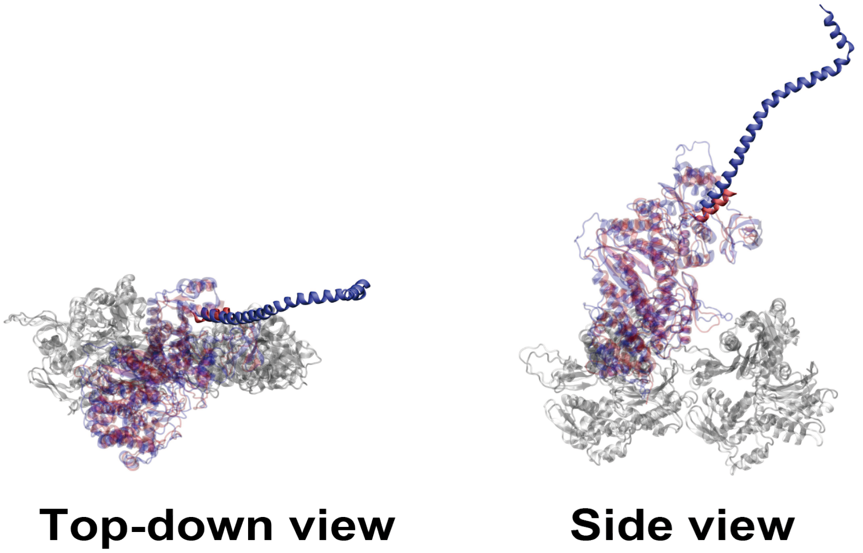


**Fig. S6. Comparison of lever arm positions in acto-*Cc*XI MD and acto-SkII S1.**
The cryo-electron microscopy structures of acto-*Cc*XI MD (PDB ID: 7KCH, blue) (7) and acto-SkII S1 (PDB ID: 5H53, red) (8) were aligned at their motor domains to compare the positions of their lever arms. (Left) Top-down view along the filament axis, highlighting the relative positions of the lever arms after the power stroke. In this view, the lever arm position of acto-*Cc*XI MD in the rigor state appears nearly identical in orientation and position to that of acto-SkII S1, with both aligned almost parallel to the filament axis. (Right) Side view illustrating the spatial orientation of the lever arms after the power stroke.

**Movie S1-S8 (separate file).**

**Legends for Movies S1 to S8**

**Movie S1. Chiral curved motion of actin filaments driven by *Cc*XI MD.** This movie shows the chiral curved motion of actin filaments driven by *Cc*XI MD in the standard *in vitro* motility assay. Actin filaments, fluorescently labeled with rhodamine-phalloidin, move in the counterclockwise (CCW) direction when viewed from the actin side, which corresponds to clockwise (CW) movement when viewed from the objective lens side. The movie is shown in real time.

**Movie S2. CCW rotation of ACRs driven by *Cc*XI MD.** This movie shows actin chiral rings (ACRs) formed in the modified in vitro motility assay. ACRs, generated through collective motion of actin filaments, rotate in the CCW direction when viewed from the actin side, corresponding to CW rotation when viewed from the objective lens side. ACRs remain stably rotating at their formation sites. Actin filaments are fluorescently labeled with rhodamine-phalloidin. The movie is shown at 10× speed.

**Movie S3. Dynamic process of ACR formation by *Cc*XI MD.** This movie captures the dynamic formation process of ACRs following the introduction of ATP-containing assay buffer. Initially, actin filaments exhibit parallel, stream-like flow, which gradually coalesces into nascent ring-like structures and eventually forms fully developed ACRs. The resulting ACRs rotate in the CCW direction when viewed from the actin side. Actin filaments are labeled with Cy3. The movie is shown at 40× speed.

**Movie S4. Effect of villin on ACR formation.** This movie shows the accelerated formation of small ACRs in the presence of 1.5 μM villin. ACRs form within one minute—significantly faster than in the absence of villin—and exhibit stable CCW rotation at their formation sites. The outer diameter of these ACRs is approximately 5 μm, which is markedly smaller than that of ACRs formed without villin. Actin filaments are fluorescently labeled with rhodamine-phalloidin. The movie is shown at 20× speed.

**Movie S5. Effect of methylcellulose on ACR formation.** This movie shows the formation of large, vortex-like ACRs in the presence of 0.5% methylcellulose. These ACRs rotate in the CCW direction when viewed from the actin side but are unstable and rapidly disassemble after formation. Actin filaments are fluorescently labeled with rhodamine-phalloidin. The movie is shown at 20× speed.

**Movie S6. Combined effects of villin and methylcellulose on ACR formation.** This movie shows the combined effects of 1.5 μM villin and 0.5% methylcellulose on ACR formation. The resulting ACRs are slightly larger in diameter (~30 μm) than those formed with villin alone. However, similar to the vortex-like ACRs formed in the presence of methylcellulose, these structures are less stable and prone to disintegration. The movie is shown at 20× speed.

**Movie S7. Simulation of collective motion of straight actin filaments.** This movie illustrates a simulation of collective motion of straight actin filaments forming nematic streams. The filaments align along their longitudinal axes via nematic interactions, resulting in bidirectional streaming. This behavior reflects the fundamental characteristics of nematic alignment in straight filament systems.

**Movie S8. Simulation of collective motion of chiral curved actin filaments.** This movie illustrates a simulation of collective motion of chiral curved actin filaments, leading to the formation of ACRs. The simulation replicates experimental observations and demonstrates that the collective motion of filaments with chiral curved motion results in the formation of ACRs.
